# Water droplet-in-oil digestion method for single-cell proteomics

**DOI:** 10.1101/2021.12.13.472378

**Authors:** Takeshi Masuda, Yuma Inamori, Arisu Furukawa, Kazuki Momosaki, Chih-Hsiang Chang, Daiki Kobayashi, Hiroto Ohguchi, Yawara Kawano, Shingo Ito, Norie Araki, Shao-En Ong, Sumio Ohtsuki

**Author notes:** **Corresponding author: Takeshi Masuda, Ph.D.** Department of Pharmaceutical Microbiology, Faculty of Life Sciences, Kumamoto University, 5-1 Oe-honmachi, Chuo-ku, Kumamoto 862-0973, Japan.

## Abstract

Recent advances in single-cell proteomics highlight the promise of sensitive analyses in limited cell populations. However, technical challenges remain for sample recovery, throughput, and versatility. Here, we first report a water droplet-in-oil digestion (WinO) method based on carboxyl-coated beads and phase transfer surfactants for proteomic analysis using limited sample amounts. This method was developed to minimize the contact area between the sample solution and the container to reduce the loss of proteins and peptides by adsorption. This method increased protein and peptide recovery 10-fold as well as the number of quantified transmembrane proteins compared to an in-solution digestion (ISD) method. The proteome profiles obtained from 100 cells using the WinO method highly correlated with those from 10000 cells using the ISD method. We successfully applied the WinO method to single-cell proteomics and quantified 462 proteins. Using the WinO method, samples can be easily prepared in a multi-well plate, making it a widely applicable and suitable method for single-cell proteomics.

## Introduction

In recent decades, single-cell omics has become an important analytical technique in several research fields that has brought new perspectives to cancer genomics^1-3^, tissue development^4^, and cellular differentiation ^5-8^. The genome and transcriptome are currently the main targets of single-cell omics studies. Quantitative amplification and next-generation sequencing enable high-throughput single-cell epigenetic and transcriptional analyses. Proteins are important biomolecules playing a major role in biological phenomena. Furthermore, because protein expression levels are reportedly difficult to predict based solely on mRNA expression levels ^9,10^, there remains a need to measure protein expression directly with proteomics.

For single-cell proteomics, high recovery of proteins and peptides, as well as high throughput, are required. To quantify proteins by proteomics, extracted proteins are digested into peptides by enzymes and then analyzed by nano-liquid chromatography-tandem mass spectrometry (nanoLC-MS/MS). Additionally, no current method can amplify proteins. Hence, it is critical to reduce adsorption losses during sample preparation and to enhance protein extraction and digestion in single-cell proteomics. Several sample preparation methods, such as single-cell proteomics by mass spectrometry (SCoPE-MS) ^11^, nanodroplet processing in one-pot for trace samples (nanoPOTS) ^12,13^, and surfactant-assisted one-pot sample preparation coupled with mass spectrometry (SOP-MS) ^14^, can dramatically improve the sample recovery rate and sensitivity of MS for single-cell proteomics. Using these approaches combined with state-of-the-art LC-MS systems, the number of proteins identified from single cell was dramatically increased. SCoPE-MS is based on multiplexing with a tandem mass tag (TMT) reagent where small amounts of samples are mixed with a carrier containing large amounts of peptides, thus reducing sample loss during LC injection. In addition, the greater signal intensity of peptides from the carrier proteome can increase to the number of MS/MS triggers. However, because the SCoPE-MS uses an in-solution digestion (ISD) method, there will be loss of proteins and peptides at the stage of sample preparation prior to LC-MS analyses^11^. The nanoPOTS method uses a specially fabricated nano-well chip and a liquid handling system for digestion. These devices were designed to process the sample in a small volume to reduce protein and peptide adsorption loss ^12,13^. However, because these devices are not yet commercially available, and the workflow is tied to the microfluidic system, the versatility of the nanoPOTS is lower than that of other approaches. The SOP-MS was developed for label-free single-cell proteomics ^14^. This approach eliminates all sample transfer steps. The sample preparation was performed in the presence of an MS-compatible surfactant ^15^, n-dodecyl-β-D-maltoside, to reduce the adsorption loss of the samples by blocking protein adsorption on plastic surfaces ^14^. The SOP-MS was successfully applied to sorted single cells and small tissue sections obtained by laser microdissection. However, the throughput of this approach on the nanoLC-MS/MS is limited due to the lack of a multiplexing approach. While these advances enable single-cell proteomics, technical challenges remain for the recovery rate, versatile application, and increased throughput.

Herein, we report a simple and highly efficient sample preparation method for single-cell proteomics that prepares samples in a water droplet, terms water droplet-in-oil digestion (WinO). This new method reduces sample loss during single-cell protein preparations and increases the number of identified proteins compared with the ISD methods. The WinO method improves current single-cell proteomics methods and can enhance the throughput and protein identification from single-cell sampling.

## Methods

### Reagents and chemicals

Sodium deoxycholate (SDC), sodium lauryl sarcosinate (SLS), ammonium bicarbonate (AmBic), dithiothreitol (DTT), iodoacetamide (IAA), mass-spectrometry-grade lysyl endopeptidase (Lys-C), ethyl acetate, acetonitrile, acetic acid, trifluoroacetic acid (TFA), Dulbecco’s modified Eagle medium (DMEM), negative staining kit, and RPMI-1640 medium were purchased from Fujifilm Wako (Osaka, Japan). Triethylammonium bicarbonate (TEAB), SOURCE 30S beads, and SP Sepharose High Performance beads were from Sigma-Aldrich (St. Louis, MO, USA). Modified trypsin was obtained from Promega (Madison, WI, USA). SDC-XC StageTip was purchased from GL Sciences (Tokyo, Japan). Carboxyl-coated Magnosphere beads were obtained from JSR Life Sciences (Tsukuba, Japan). Benzonase nuclease was purchased from Merck Millipore (Burlington, MA, USA). LDS sample buffer, BCA assay kit, Tandem Mass Tags reagents, and Dynabeads MyOne Carboxylic acid were obtained from Thermo Scientific (San Jose, CA, USA). FG beads COOH and FG beads NH2 were from TAMAGAWA SEIKI (Nagano, Japan).

### Cell culture and cell sorting

In this study, 15 multiple myeloma cell lines (H929, KMM-1, KMS-11, KMS-12BM, KMS-12PE, KMS-20, KMS-27, KMS-28BM, KMS-28PE, L363, MM.1S, MOLP8, OPM1, RPMI8226, and U266 cells) were cultured in RPMI-1640 medium supplemented with 10% FCS to 80% confluence. Cells were washed three times with PBS, and then 1 or 100 cells were sorted into 96-well plates. For 10000 cells, cells were sorted into 1.5 mL tubes. As a cell sorter, an SH800S Cell Sorter (Sony, Tokyo, Japan) using a 100-μm chip was used. Dead and doublet cells were removed prior to cell sorting. HEK293 cell was cultured in DMEM supplemented with 10% FCS to 80% confluence. Cells were harvested by scraper and washed three times with PBS.

### Examination of proteins and peptides retention in water droplet in ethyl acetate

The protein solution was prepared by extraction from HEK293 cells using the 12 mM SDC and 12 mM SLS in 100 mM Tris-HCl (pH 9.0). The protein amount was quantified by the BCA assay kit. To examine protein retention in water droplets in ethyl acetate, 50 μL of solution containing 10 μg of HEK293 proteins was dropped into 500 μL of ethyl acetate in a 1.5 mL tube. The samples were incubated for 24 hours at 25 °C. The ethyl acetate and water droplet were transferred by pipette tip into a new 1.5 mL tube and evaporated with a centrifuge concentrator. Each fraction was reconstituted with 20 μL of LDS sample buffer and separated by 5-20% SDS-PAGE. To prepare the negative control sample, we performed the same process as described above using the extraction buffer containing 12 mM SDC and 12 mM SLS in 100 mM Tris-HCl (pH 9.0). As the positive control, 10 μg of HEK293 proteins were separated by SDS-PAGE together. The protein bands in the gel were detected by the negative staining.

To examine the peptide retention in the water droplet in ethyl acetate, the HEK293 peptides prepared by the ISD method were used. To prepare the peptide solution, 10 μg of HEK293 proteins were reduced and alkylated with 10 mM DTT and 50 mM IAA, respectively. The protein solution was diluted 4-fold with 50 mM AmBic prior to enzymatic digestion. Proteins were digested with 0.5 μg of Lys-C followed by 0.5 μg of trypsin overnight at 37 °C. For the evaluation, 20 μL of solution containing 10 μg of HEK293 peptides was dropped into 200 μL of ethyl acetate and incubated at 25 °C for 24 hours. The ethyl acetate and water droplet were transferred by pipette tip into a new 1.5 mL tube and dried with the centrifuge concentrator. Each fraction was reconstituted by the 50 μL of 50 mM AmBic and subjected to the phase transfer method to remove SDC and SLS. The peptides purified with SDB-XC StageTip were analyzed by nanoLC-MS/MS using TripleTOF 5600 (Sciex, Framingham, MA). As the positive control, 10 μg of HEK293 peptides purified by SDB-XC StageTip were used for nanoLC-MS/MS. To prepare the negative control sample, we performed the same process as described above using the buffer containing 3 mM SDC, 3 mM SLS, 37.5 mM AmBic in 25 mM Tris-HCl (pH 9.0).

### Sample preparation using the ISD method

First, 1 μL extraction buffer (50 mM TEAB, 12 mM SDC, 12 mM SLS, and 0.125 units benzonase) was added into low protein-binding 96-well plates (Sumitomo Bakelite, Tokyo, Japan) or into low protein-binding 1.5 mL tubes (Watson, Tokyo, Japan). Cells were sorted into each well and spun at 300 × *g* for 1 min at 25 °C to mix the extraction buffer and the cells. The reduction and alkylation were performed by adding 1 μL 100 mM DTT and 550 mM IAA solution, respectively. The DTT and IAA solutions were prepared by 50 mM TEAB. Proteins were digested with Lys-C for 3 hours at 37 °C, followed by trypsin incubation for 16 hours at 37 °C. Enzymes were prepared with 50 mM TEAB, and 50 ng and 0.5 ng of enzymes were used to digest for 100 cells and a single cell, respectively (1 μL solution was used). The plates were sealed with an adhesive plate seal for every incubation. After digestion, 4 μL (40 μg) TMT reagent in 0.5% acetic acid and 50% acetonitrile were added, and samples were incubated for 60 min at 25 °C. The pH of the sample solutions during TMT labeling was approximately pH 8. To quench the TMT reaction, 1 μL 30% hydroxylamine was added to each sample and incubated for 15 min at 25 °C. Subsequently, the surfactants were removed using the phase transfer method ^16,17^. Briefly, sample solutions, including ethyl acetate, were combined and acidified with TFA to give a final concentration of 0.5%. The combined samples were mixed by vortexing and then centrifuged at 15600 × *g* for 2 min. Ethyl acetate, including surfactants, was discarded. The peptides were purified using SDB-XC StageTip ^18,19^.

### Sample preparation using the WinO method

For the WinO digestion, 50 μL ethyl acetate were added into the wells of a 96-well plate before adding 1 μL extraction buffer (50 mM TEAB, 12 mM SDC, 12 mM SLS, and 0.125 units of benzonase). Cells were sorted into each well and spun at 300 × *g* for 1 min at 25 °C to mix the extraction buffer droplets and cell droplets. After 30 min incubation at RT, 1 μL (3.3 μg) carboxyl-coated Magnosphere beads equilibrated with 50 mM TEAB were added to each well. From the reduction step with 100 mM DTT, the sample preparation was performed as per the ISD method. The solutions were added into ethyl acetate.

### nanoLC-MS/MS analysis

Three different mass-spectrometry systems were used in this study. To determine the optimum magnetic beads for the WinO method, a TripleTOF 5600 (Sciex, Framingham, MA) and an UltiMate 3000 RSLCnano (Thermo Fisher Scientific) were used. In this system, 5 μL was injected into the LC system. The LC was performed using an Acclaim PepMap RSLC (75 μm × 25 cm, C18, 2 μm, Thermo Fisher Scientific) at 300 nL/min. The mobile phase consisted of (A) 0.1% formic acid and (B) 0.1% formic acid in acetonitrile. Linear gradients of 2–25% B in 60 min, 50–90% B in 15 min, and 90% B for 5 min were applied, and the spray voltage was 2300 V. For information-dependent acquisition (IDA), the precursor scan range was *m/z* 300– 1250 in 250 ms. The top 20 precursor ions with a charge of +2 to +5 were selected, and the product ion scan was performed for 50 ms over the range of *m/z* 100–1600. Sequential window acquisition of all theoretical fragment-ion spectra (SWATH) was performed with the LC method described for IDA using 73 variable windows with 40 ms scan times. The product ions were collected in the range of *m/z* 100–1600 in high-sensitivity mode.

The 100-cell proteomic analysis utilizing the TMT reagents was performed on an Orbitrap Fusion Tribrid mass spectrometer (Thermo Scientific) and an EASY-nLC™ 1200 system (Thermo Scientific). Peptides were first loaded onto an Acclaim™ PepMap™ 100 C18 (3 μm, 75 μm ID× 20 mm length, P/N 164946 Thermo Scientific), and then separated on C18 packed emitter column (3 μm, 75 μm I.D. × 150 mm length, Nikkyo Technos, Tokyo, Japan). The injection volume was 5 μL and the flow rate was 300 nL/min. The mobile phase consisted of (A) 0.1% formic acid and (B) 0.1% formic acid in 80% acetonitrile. A multiple-linear gradient elution was performed as follows: 5–35% B in 60 min, 35–100% B in 5 min, and 90% B for 10 min. Multiplex analysis using TMT reagents was performed using a synchronous precursor selection (SPS) MS3 scan in top-speed mode (cycle time = 3 sec). The parameters were as follows: spray voltage, 2300 V; temperature of the ion transfer tube, 250 °C; Orbitrap scan range for precursor ion (*m/z*), 350–1500; resolution for precursor scan, 120,000; ion trap scan range (*m/z*), auto; collision energy for MS2, 35%, collision mode for MS2, CID; collision energy for MS3, 65%; collision mode for MS3, HCD; Orbitrap scan range for MS3 (*m/z*), 100–500; resolution for reporter ion detection, 50,000; maximum injection time, 50 ms for MS1 and MS2 scan,105 ms for MS3 scan; AGC target, 400,000 for MS1 orbitrap, 10,000 for MS2 ion trap, 100,000 for MS3 orbitrap.

For single-cell proteomics utilizing the TMT reagents, we performed the SPS MS3 scan with a real-time search based on an Orbitrap Eclipse Tribrid mass spectrometer (Thermo Scientific) with FAIMS Pro interface (Thermo Scientific) and an EASY-nLC™ 1200 system (Thermo Scientific) equipped with an Aurora column (75 μm I.D., 15 cm length, 1.6 μm beads, IonOpticks, Fitzroy, Australia). The mobile phase consisted of (A) 0.1% formic acid and (B) 0.1% formic acid in 80% acetonitrile. LC gradients of 6–20% B in 37 min, 20–30% B in 14 min, 30–40% B in 9 min, 30–90% B in 3 min, and 90% B for 7 min were applied. For the SPS MS3 scan, the parameters were as follows: flow rate, 300 nL/min; spray voltage, 2000 V; temperature of ion transfer tube, 275 °C; Orbitrap scan range for precursor ion (*m/z*), 375–1500; resolution for precursor scan, 60,000; FAIMS CV, -50 and -70; ion trap scan range (*m/z*), 200–1200; collision energy for MS2, 30%; collision mode for MS2, CID; injection time for MS2 (ms), 500; collision energy for MS3, 65%; collision mode for MS3, HCD; Orbitrap scan range for MS3 (*m/z*), 100–500; reporter ion detection resolution, 50,000; injection time for MS3 (ms), 500; AGC target, 200% (1E5). The parameters for the real-time search were as follows: enzyme, trypsin; maximum search time (ms), 100; SPS mode, true; Xcorr, 1.4; sCn, 0.1; precursor, 10 ppm; precursor range (*m/z*), 400–1200; precursor exclusion, low 25 ppm and high 25 ppm.

### Data analysis

The IDA and SWATH data obtained by the TripleTOF 5600 were analyzed using ProteinPilot version 4.5 (Sciex) and PeakView version 2.1 (Sciex). The false discovery rate (FDR) was estimated by searching against a decoy database generated by randomization of the UniProt human reference database. The data obtained from the Orbitrap Fusion Tribrid mass spectrometer was analyzed using MaxQuant ^20^ ver.1.6.17.0, and the UniProt human reference database was used as the reference. The parameters were as follows: type of parameter section, reporter ion MS3; isobaric labels, 10plex TMT or 11plex TMT; enzyme, trypsin/P; variable modifications, oxidation (M) and acetyl (protein N-term); and fixed modifications, carbamidomethyl (C). For the data obtained from the Orbitrap Eclipse Tribrid mass spectrometer, we used Proteome Discoverer ver.2.5 (Thermo). The parameters were as follows: enzyme, trypsin (full); maximum missed cleavage site, 2; and dynamic modification, oxidation (M), acetyl (N-terminal of protein), Met-loss (N-terminal of protein), and Met-loss+ acetyl (N-terminal of protein); static modification, carbamidomethyl (C); minimum average reporter S/N, 10. According to a previous report, the grand average of hydropathy (GRAVY) scores of proteins and peptides were calculated ^21^. For gene ontology (GO) analysis, we used DAVID version 6.7 (https://david.ncifcrf.gov/summary.jsp). Uniform manifold approximation and projection (UMAP) was generated using the R script. The *t*-test was performed using GraphPad Prism 8.4.3 (GraphPad Software, CA) or Excel (Microsoft, MA). One-way ANOVA Tukey’s multiple comparison test was performed using GraphPad Prism 8.4.3. Statistical significance in the proteomics data was considered for fold changes ≥ 2 or ≤ 0.5 with a *p*-value < 0.05.

## Results and discussion

### 1. Effect of the WinO method on the recovery of small samples

In the WinO method, a 1-μL water droplet containing 50 mM TEAB, 12 mM SDC, 12 mM SLS, and 0.125 units benzonase was formed in 50 μL ethyl acetate (Figure 1A). The SDC and SLS have been reported to enhance protein extraction and digestion efficiencies of Lys-C and trypsin ^16,17^. In addition, SDC and SLS are known as phase transfer surfactants (PTSs), which can be removed from peptide solutions by a phase transfer method ^16,17^. To enhance the protein extraction from cells, heating, ultrasonication, and freezing/thawing are generally used. In the WinO method, these treatments are not possible because ethyl acetate is volatile, or sample solution and ethyl acetate are mixed during these treatments. Hence, we assessed differences in extraction efficiency between ultrasonication and heating in the PTS solution and a solution prepared by mixing cells in the PTS solution (Figure S1). The protein amount, number, and intensity of quantified proteins and peptides were comparable between the two methods. Therefore, ultrasonication and heat treatment were not used for the protein extraction process in this study. The magnetic beads, DTT, IAA, Lys-C, and trypsin solutions prepared in 50 mM TEAB were added to ethyl acetate. These solutions form water droplets in ethyl acetate, which merge with the sample droplet. The digested peptides were labeled by adding the TMT solution; after combining multiple samples, SDC and SLS were removed using the phase transfer method ^16,17^ (Figure 1B). The peptides purified with StageTip were analyzed by nanoLC-MS/MS.

**Figure 1.**
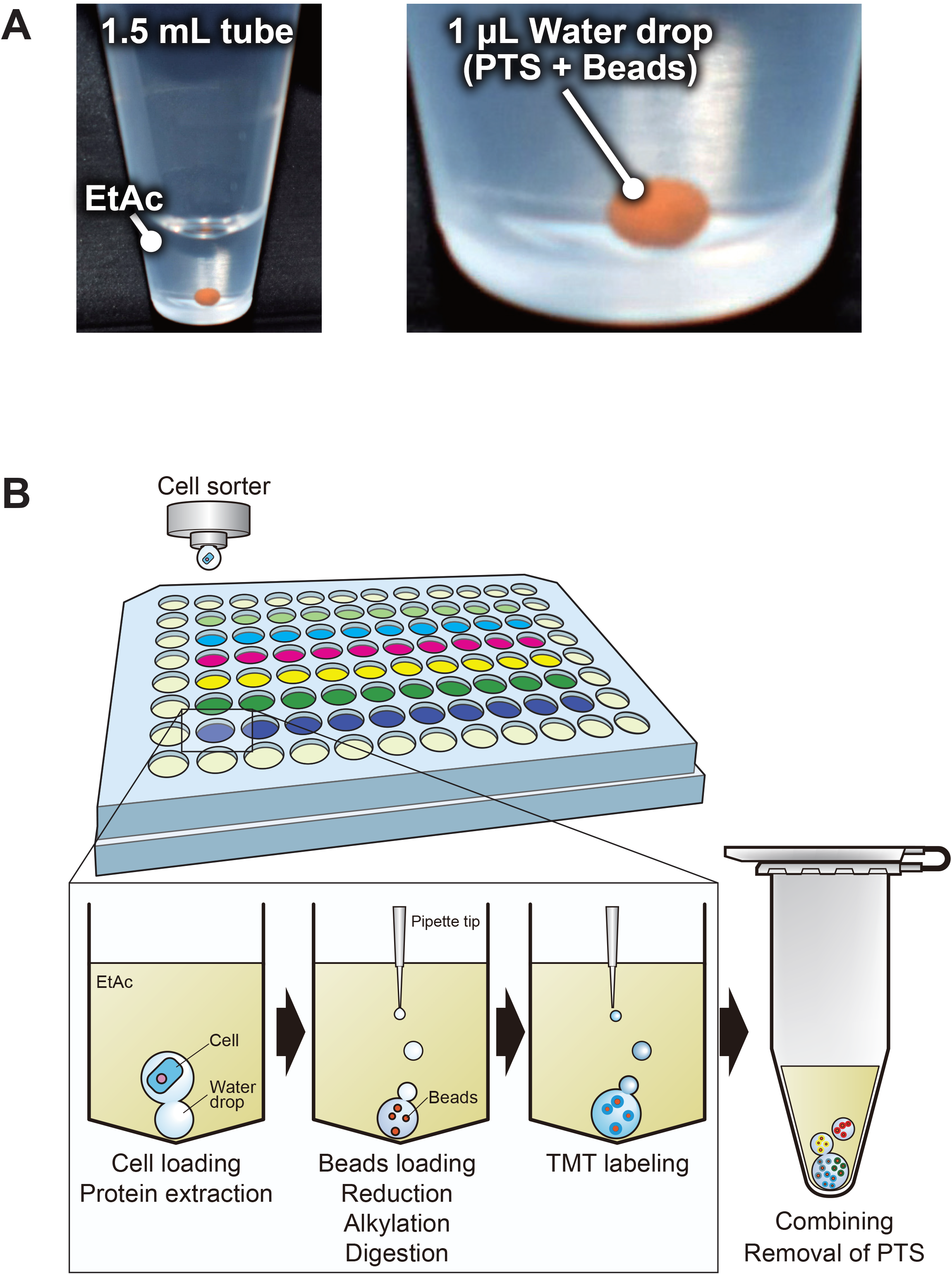
Water droplet-in-oil digestion (WinO) method. One microliter of water droplet containing 0.125 units of benzonase, 3.3 μg magnetic beads, 12 mM sodium deoxycholate (SDC), 12 mM sodium lauroyl sarcosinate (SLS), and 100 mM TEAB (pH 8.5) formed in ethyl acetate (A). The workflow of the WinO method is presented in (B); the cells are loaded into water droplets in ethyl acetate using a cell sorter. The solutions for reduction, alkylation, and digestion are added to the ethyl acetate. The peptides are labeled with TMT reagents and then combined. The peptides are purified using the StageTip and injected into the nanoLC-MS/MS.

To examine whether the proteins and peptides were retained in the water droplet in ethyl acetate, 10 μg HEK293 whole cell lysate or 10 μg digested peptides solution were added into ethyl acetate and incubated for 24 h. Ethyl acetate and water droplets were collected, and the distribution of each fraction was confirmed by SDS-PAGE and nanoLC-MS/MS for proteins and peptides, respectively. As controls, whole-cell lysates or peptide solutions were also used for SDS-PAGE or nanoLC-MS/MS along with the treated samples. In addition, unloaded samples without proteins or peptides were prepared as negative control samples. To examine protein retention in water droplets in ethyl acetate using SDS-PAGE, smears of proteins larger than 100 kDa and smaller than 25 kDa were detected in the ethyl acetate fraction (Figure S2A). These smears were also detected in the ethyl acetate fraction of the unloaded negative control samples. No other protein bands were detected in the ethyl acetate fraction of the protein-loaded group (Figure S2A). Next, we examined the distribution of the peptides in ethyl acetate and the sample droplet.

There was no significant difference in the total peak area of the peptides detected in the control and sample droplet fractions (*p* = 0.5734, Figure S2B). The percent composition of the peptide peak area quantified in the ethyl acetate fraction was only 0.12%. The total peak area of peptides in this fraction showed no significant difference from that in the ethyl acetate fraction of negative control (*p* = 0.0561) (Figure S2B). These results suggest that proteins and peptides are retained in water droplets for at least 24 hours.

Next, we performed the WinO method using sorted cells as starting material, which was then compared to the recovery of peptides (Table S1) and proteins (Table S2) in the ISD method. The WinO method was performed as described above. In the ISD method, surfactant solution was added into the well of a 96-well plate before 100 cells were injected into the solution using a cell sorter. Next, the DTT, IAA, Lys-C, and trypsin solutions were added directly into the sample solution. TMT-labeled peptides corresponding to 5000 cells as carriers were combined with the 100-cell samples from the ISD and WinO methods. The carrier was used to increase peptide and protein identification, as well as reduce peptide loss after digestion as SCoPE-MS ^11^. The peptide intensity in the WinO method was significantly higher than that in the ISD method (*p* < 0.0001; n = 3, Figure 2A). Next, we compared the recovery of the peptides quantified in all data. The intensity of 1018 out of 1177 peptides (86.5%) increased significantly (≥ 2-fold, *p* < 0.05) in the WinO method, whereas the intensity of none of the peptides increased significantly in the ISD method (Figure 2B). The median relative peptide recovery from WinO was 6.70-fold greater than that from the ISD method. The number of quantified peptides increased only 1.3-fold (*p* = 0.0199) when the WinO method (2071.7 ± 34.4) was used compared to the ISD method (1598.3 ± 215.9). In this study, we counted the proteins in which reporter ions were detected. The triggering of MS2 was assisted by the carrier, which reduced the difference in the number of detected proteins between the two methods. To examine the reproducibility of the WinO method, we compared the % coefficient of variations (CVs) of peptide levels in triplicate between the two methods. The %CV distribution pattern using the WinO method was lower than that with the ISD method (Figure 2C), with median %CVs of 39.6% and 14.4% for ISD and WinO methods, respectively.

**Figure 2.**
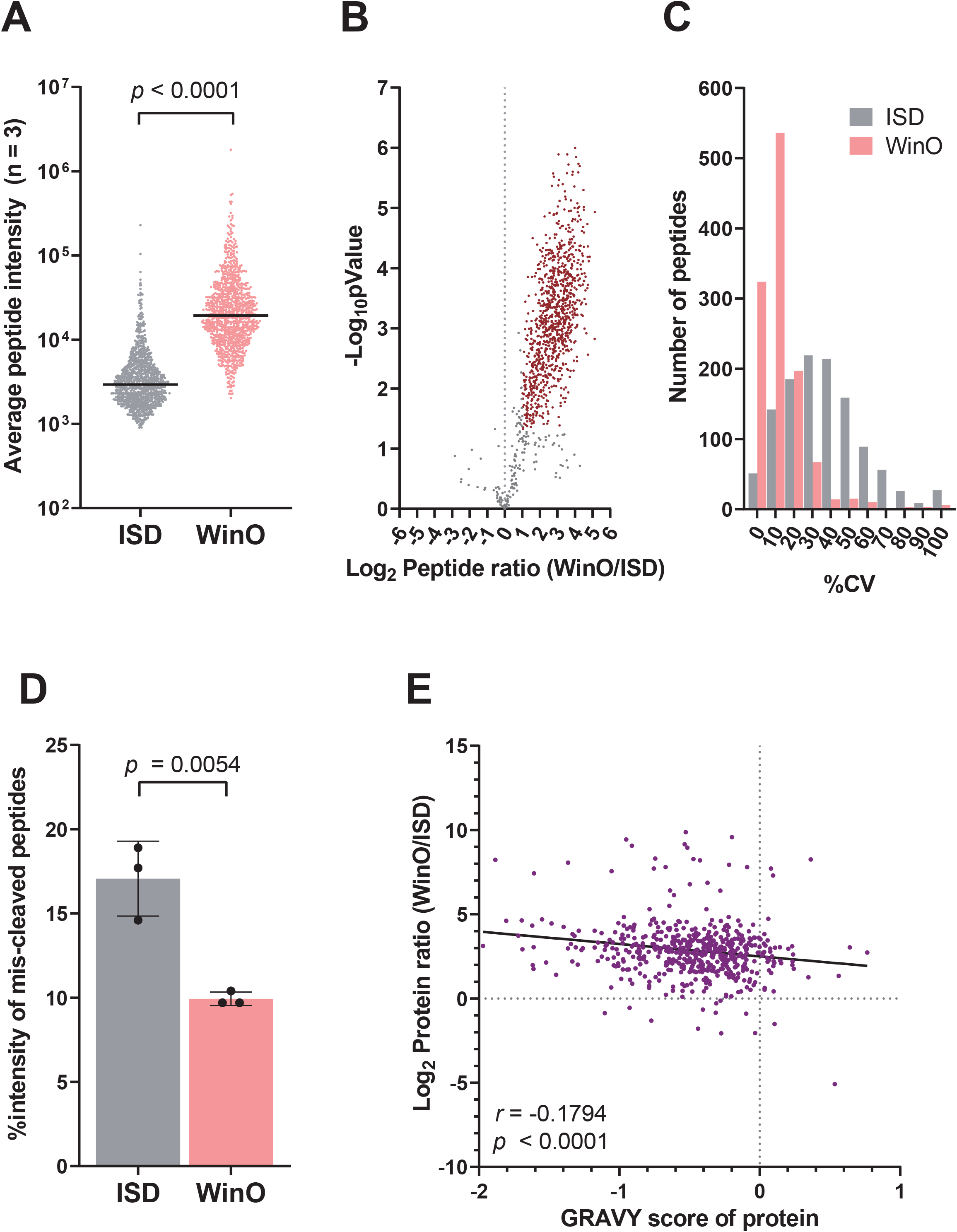
Comparison of the in-solution digestion (ISD) and WinO methods. As starting material, 100 RPMI8226 cells were sorted and digested in triplicate using ISD and WinO methods. The scatter plot shows the levels of 1015 peptides quantified using these digestion methods (A). Peptide levels are represented as the average of triplicate data. Each bar shows the median. Paired *t*-test was performed using GraphPad Prism 8.4.3. The relative peptide levels between the WinO and ISD methods are shown as a volcano plot (B). Red dots indicate peptides with a significant change (*p* < 0.05, 2-fold or more). The distribution of %CV from the ISD and WinO methods is presented in (C). The %CV for peptide intensities was calculated from triplicate data generated using each method. The proportion of mis-cleaved peptides in the ISD and WinO methods is shown in (D). These proportions were calculated based on peptide levels and averaged across triplicate samples. Error bars indicate standard deviation. Unpaired *t*-test was performed using GraphPad Prism 8.4.3. The correlation between the GRAVY protein score and relative protein levels from the WinO to the ISD method is shown in (E). The GRAVY score of proteins was calculated as previously reported ^21^. In this correlation, 561 proteins commonly quantified by both methods are shown. The Pearson correlation and *p*-values were calculated using GraphPad Prism 8.4.3.

Next, we evaluated the digestion efficiency of the WinO method by measuring the levels of mis-cleaved peptides. Lys-C and trypsin solutions were delivered to the sample droplets through ethyl acetate in the WinO method, and 8.03 g ethyl acetate was dissolved in 100 mL water at room temperature ^22^. It was expected that the dissolved ethyl acetate affects the activity of Lys-C and trypsin. However, mis-cleavage of the total peptides in the WinO method (9.9 ± 0.4%) was significantly lower than that in the ISD method (17.1 ± 2.2%) (Figure 2D). In addition, the improvement in peptide recovery from fully cleaved peptides was significantly higher (*p* < 0.0001) than in the mis-cleaved peptides using the WinO method (Figure S3). Contrary to expectations, the digestion efficiency of the WinO method was enhanced compared to that of the ISD method. It was reported that the activities of proteolytic enzymes are enhanced in the presence of organic solvents, such as methanol, isopropyl alcohol, and acetonitrile ^23^. It is likely that the dissolved ethyl acetate in the sample droplet enhanced Lys-C and trypsin activities in the WinO method. Moreover, the reduction in the adsorption loss of Lys-C and trypsin maintained a high enzyme concentration in the sample droplet. Although the recovery of proteins was overall improved with the WinO method, a significant negative correlation (*r* = -0.1794, *p* < 0.0001) was observed in protein hydrophobicity, as evidenced by the GRAVY score and protein recovery (Figure 2E). A greater score indicates a more hydrophobic protein/peptide in GRAVY score. This indicated that the WinO method led to a higher recovery of hydrophilic than hydrophobic proteins.

Based on these results, we speculated that the 100-cell protein and peptide recoveries were enhanced with the WinO method due to the reduced contact surface area between the sample solution and plastic tubes, as well as the improved digestion efficiency of trypsin and Lys-C. However, the improvement in hydrophobic protein recovery was lower than that in hydrophilic proteins, possibly due to lower retention of these proteins in the ethyl acetate solution than hydrophilic proteins.

### 2. Effect of carboxyl-coated magnetic beads on peptides recovery using the WinO method

To enhance the recovery of hydrophobic proteins and peptides, we tried to retain them in the sample droplet using beads. To select beads with high peptide recovery for the WinO method, we examined six different types of beads, an amine-, a methyl sulfonate-, a sulfopropyl-, and three carboxyl-coated beads. The carboxyl-coated beads tended to show a higher total intensity of proteins than other bead types. The carboxyl-coated Magnosphere beads showed the highest recovery among all beads tested (Figure S4). Single-pot, solid-phase-enhanced sample preparation (SP3) ^24^ and protein aggregation capture (PAC) ^25^ have been reported as methods to retain peptides on beads. In SP3 and PAC, peptides are captured on the beads in organic solvent based on the hydrophilic interaction. In this study, proteins and peptides were retained on the beads in aqueous solution. Hence, we assumed that proteins and peptides are retained on the beads via ionic interactions rather than hydrophilic interactions. In addition, it is likely that the hydrophobic parts of proteins and peptides have a high affinity with the beads because of the hydrophobic material of Magnosphere beads. We performed proteomic analysis of the 100 sorted cells using the WinO method with or without beads in triplicate (Tables S3 and S4 for peptides and proteins, respectively). After digestion, each sample was labeled with TMT reagent and mixed. The peptide level significantly increased in WinO samples combined with beads compared to WinO samples without beads (*p* < 0.0001; Figure 3A). Figure 3B shows the distribution of relative peptide levels (n = 1898) in the WinO samples processed with beads compared to those without. The median peptide ratio was 1.497 (Figure 3B). The peptide ratio of 1825 out of 1898 peptides (96.2%) was higher in samples prepared with the beads than without beads. Moreover, there was no significant difference in the percentage of cleaved peptides (Figure 3C), suggesting that the addition of beads does not affect the Lys-C and trypsin activities. The reproducibility of the WinO method was evaluated by comparing the protein levels from triplicate analyses (Figure 3D). The Pearson correlations for all pairs were higher than 0.96, indicating that the reproducibility of peptide quantification was unaffected by the presence of beads. Thus, we combined the beads with the WinO method in subsequent experiments.

**Figure 3.**
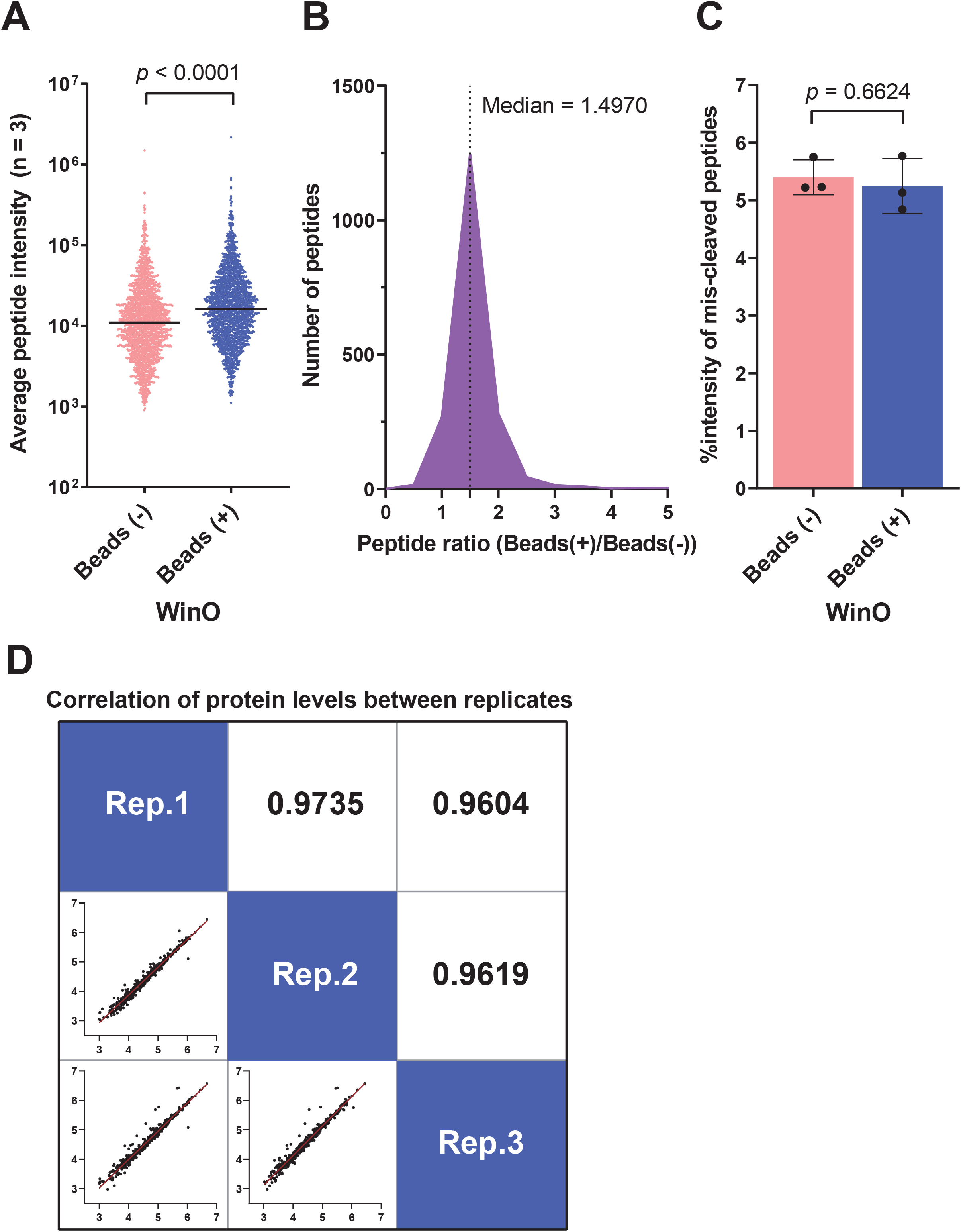
Effect of carboxyl-coated magnetic beads on the WinO method efficiency. One hundred RPMI8226 cells were sorted and digested in triplicate. The WinO method was performed with and without magnetic beads. Peptide levels are presented as the average of triplicate data. The scatter plot shows the levels of 1898 peptides quantified using both methods (A). Each bar shows the median. Paired *t*-test was performed using GraphPad Prism 8.4.3. The distribution of relative peptide levels between the WinO method with and without beads are shown in (B). The proportion of mis-cleaved peptides in the WinO method with and without beads is shown in (C). These proportions were calculated based on peptide level and averaged across triplicate samples. Error bars indicate standard deviation. Unpaired *t*-test was performed using GraphPad Prism 8.4.3. Correlation of protein levels between replicates (D). Simple linear regressions were performed using GraphPad Prism 8.4.3.

We characterized proteins and peptides whose recovery was improved by the addition of beads. Figure 4A shows the correlation of protein recovery and GRAVY score. The hydrophobicity of proteins did not correlate with their recovery; in other words, the protein recovery rate improved independently of their hydrophobicity. Next, the Spearman correlation coefficient was calculated by comparing the recovery rate of peptides with the frequency of each amino acid and GRAVY score (Figure 4B). A significant positive correlation was observed in the GRAVY score and the peptide recovery (*r* = 0.0810, *p* = 0.0004; Figure 4B). In addition, the frequency of the basic amino acids (H, K, and R) showed the highest coefficient (*r* = 0.1067, *p* < 0.0001, Figure 4B, Figure S5), whereas the frequency of acidic amino acids (D and E) showed the lowest coefficient (*r* = -0.1162, *p* < 0.0001; Figure 4B, Figure S5). The recovery of basic and hydrophobic peptides, which had a high affinity for beads in the basic condition, was improved by the WinO method. Although acidic amino acid showed a negative correlation coefficient with peptide recovery, the peptide and protein recoveries improved overall in the WinO samples combined with beads (Figure 3A, 3B, and Figure 4A).

**Figure 4.**
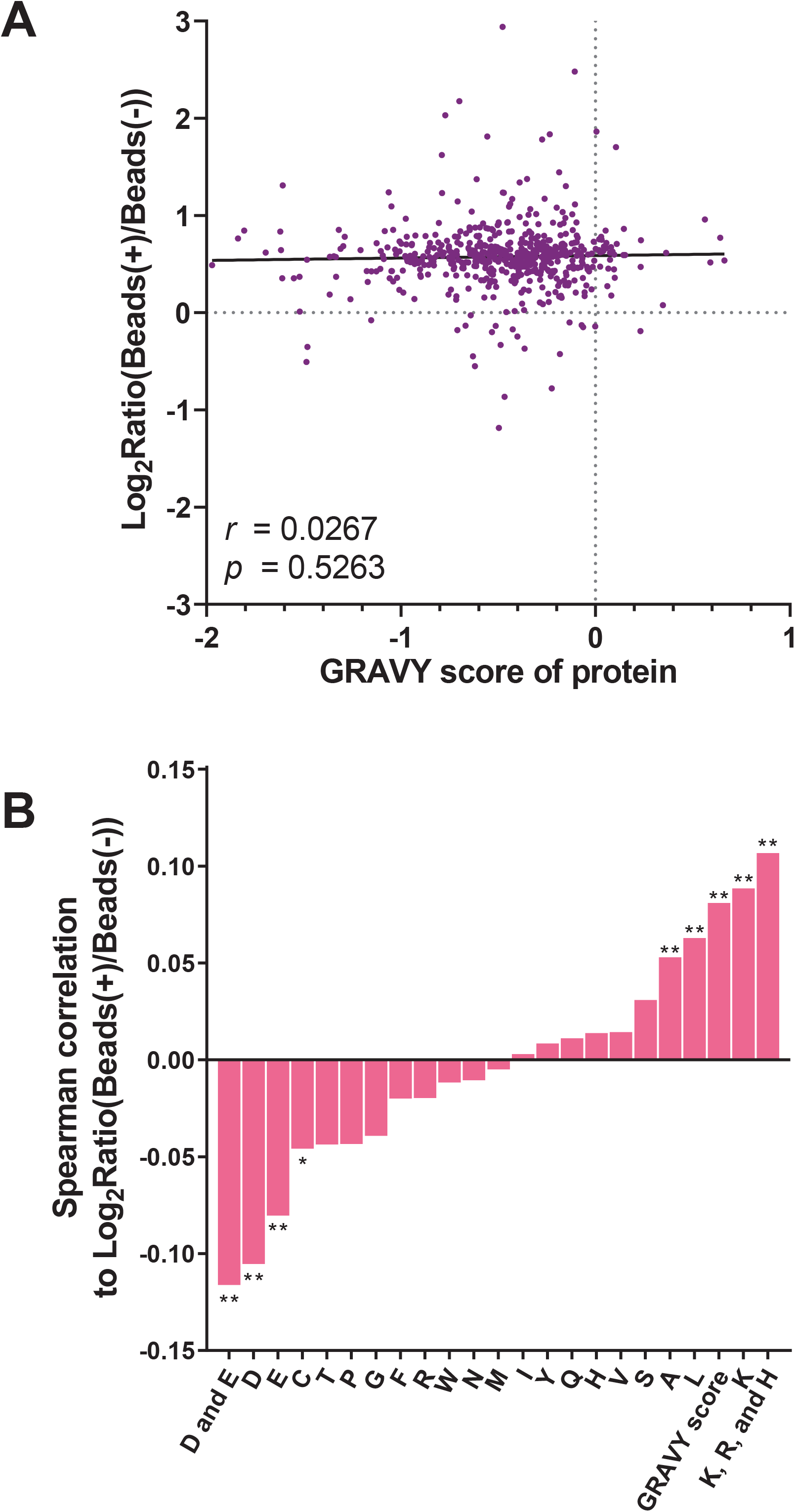
Recovery rate and characteristics of peptides using carboxyl-coated magnetic beads with the WinO method. The GRAVY score of proteins was calculated as previously reported ^21^. The Pearson correlation and *p*-values were calculated using GraphPad Prism 8.4.3 (A). The Spearman correlation coefficient was calculated by comparing the recovery rate of peptides with the WinO method with or without beads (B). The frequency of each amino acid and GRAVY score is presented for the corresponding peptides. The one-letter amino acid code is indicated on the X-axis. ** indicates *p* < 0.01; * indicates *p* < 0.05. Spearman’s correlations and *p*-values were calculated using GraphPad Prism 8.4.3.

### 3. Comparison of the proteome profiles obtained with the ISD and WinO methods

To date, none of the proteomic approaches have evaluated water droplets formed in the oil. Therefore, it was unclear whether the proteome profiles obtained by the WinO method were comparable to those obtained using the conventional ISD method. To examine the similarity of the proteome profiles between these two methods, we compared the proteome profile of 100 cells processed with the WinO method with that of 10000 cells processed with the ISD method using 15 multiple myeloma cell lines (Figure 5). The cells were sorted into a 96-well plate for the WinO method and into 1.5 mL tubes for the ISD method. From the 100-cell group, an average of 2183.6 ± 74.5 peptides were quantified (Table S5), whereas an average of 29293.0 ± 561.2 peptides were quantified from the 10000-cell group (Figure 6A, Table S6). From these peptides, an average of 592.9 ± 13.2 proteins were quantified from the 100-cell group (Table S7), whereas an average of 4651.6 ± 91.6 proteins were quantified from the 10000-cell group (Figure 6B, Table S8). In total, 798 proteins were quantified from the 15 strains using the WinO method, among which 387 proteins were found in all cell lines. Using the ISD method with 10000 cells, 5545 proteins were identified, with 3584 proteins found in all cell lines. Next, we compared the expression profiles of the 377 proteins that were quantified in both methods. Normalized expression levels were plotted using the UMAP algorithm (Figure 6C). The proteome profiles of 100 and 10000 cells were plotted close to each other and formed populations among the same cell line, even though the sample preparation method and the number of cells were different. The median %CV of the protein level for the 100-cell group (14.1%, n = 5805) was higher than that for the 10000-cell group (3.0%, n = 53760; Figure S6). The reduced reproducibility in the 100-cell group could be due to cellular heterogeneity in a limited sample, as well as higher variability in sample preparation from a small number of cells.

**Figure 5.**
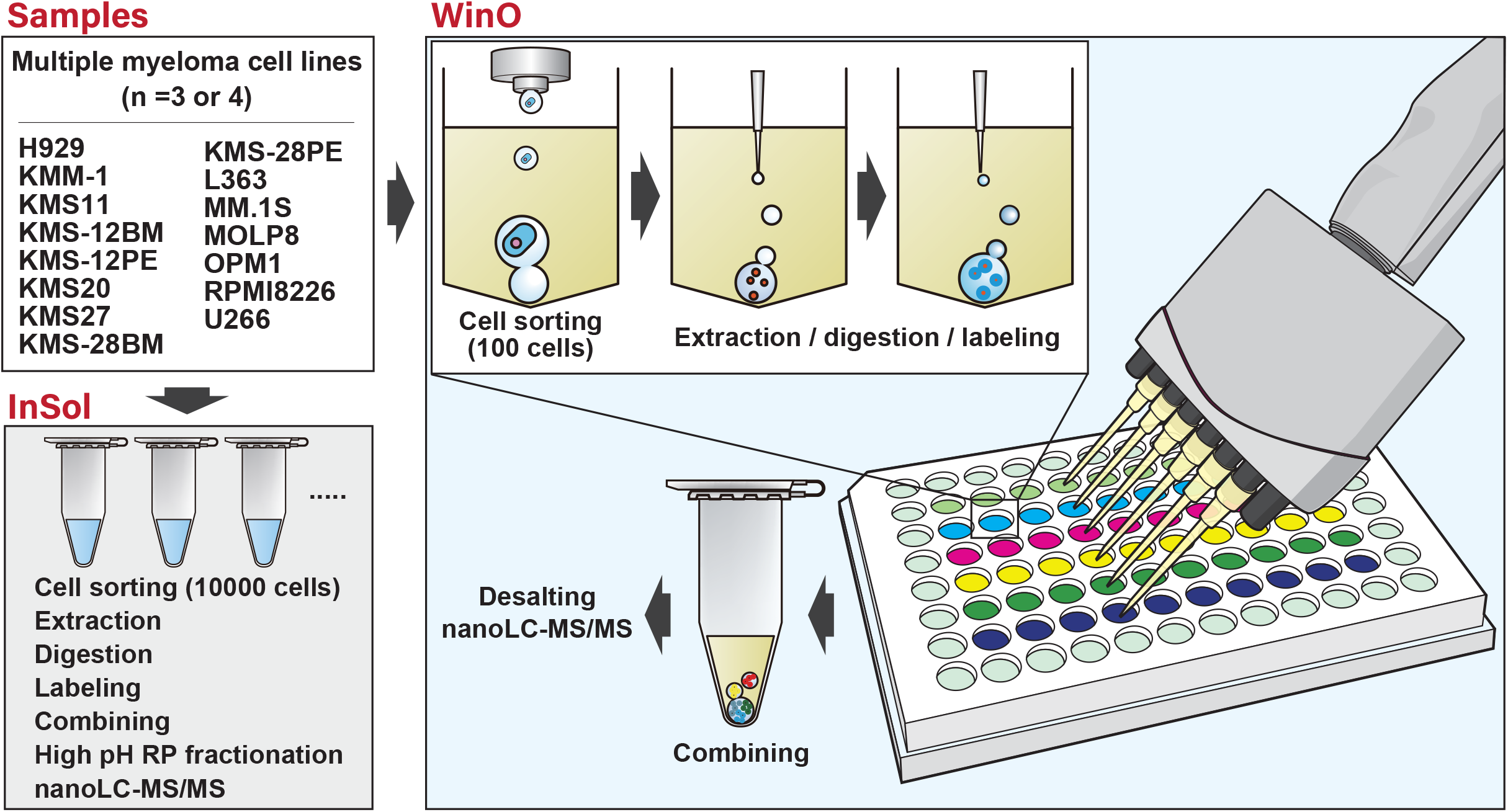
Workflow of 100-cell and 10000-cell proteomic analysis using 15 multiple myeloma cell lines. As starting materials, 10000 or 100 cells from 15 multiple myeloma cell lines were sorted and digested in triplicate or quadruplicate. For the ISD and WinO methods, 10000 or 100 cells were sorted into 1.5 mL tubes and 96-well plates, respectively. Peptides labeled with TMT reagents were combined. For the ISD method, the peptide mixture was separated into nine fractions using high-pH reverse phase fractionation.

**Figure 6.**
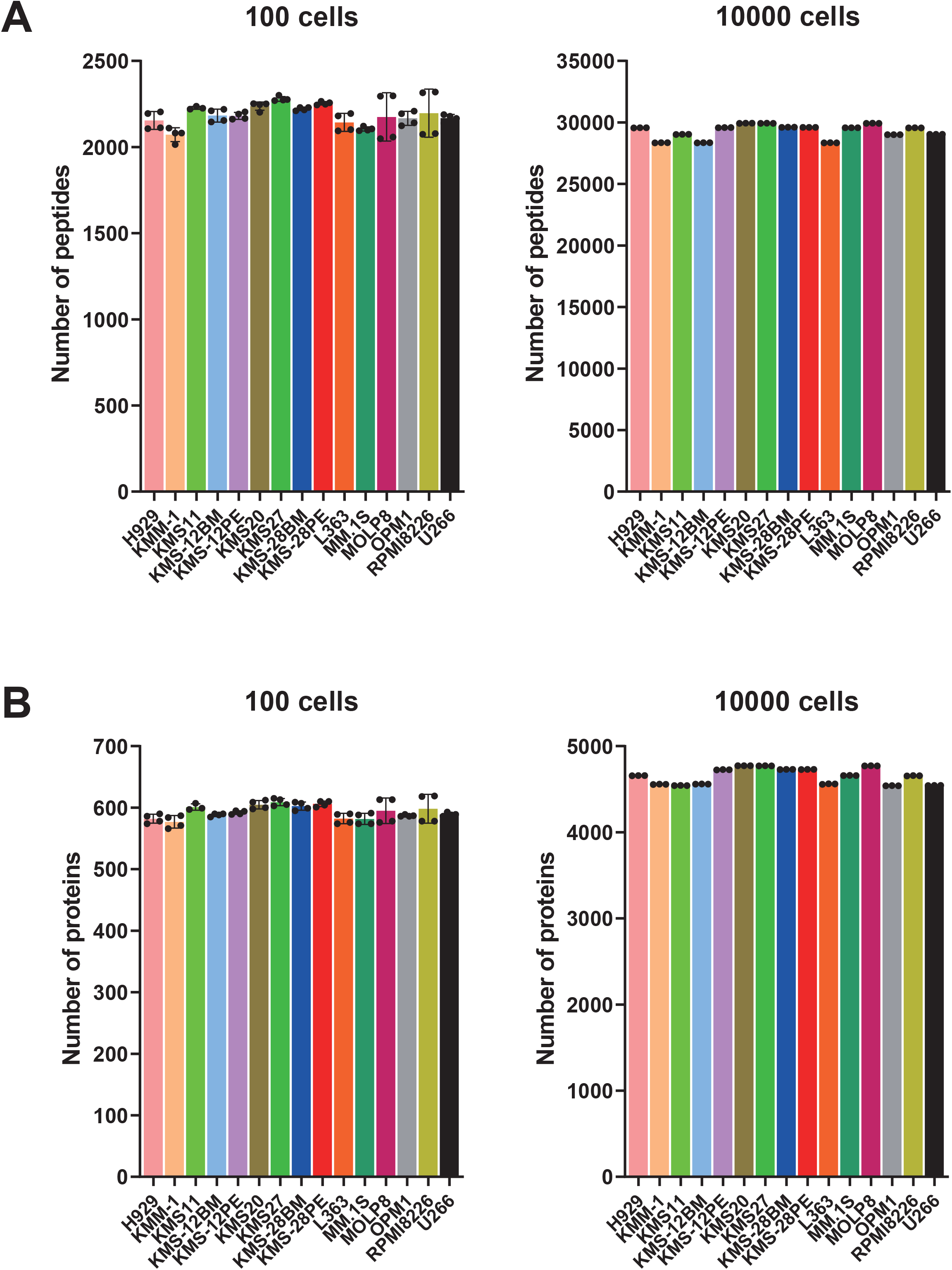
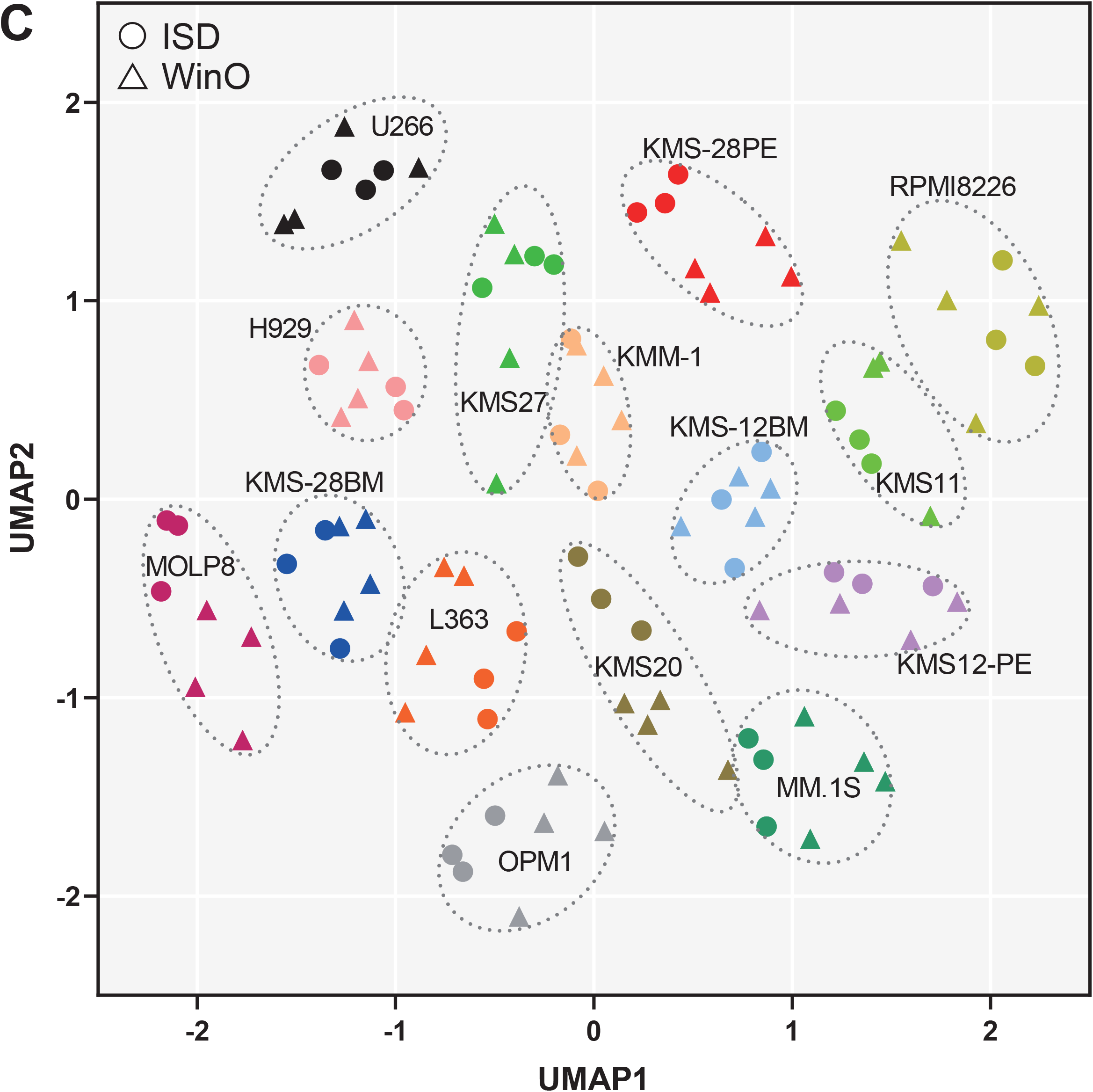
Number of peptides and proteins quantified using the ISD or WinO methods. The numbers of quantified peptides and proteins are shown in (A) and (B), respectively. Each bar indicates the average and standard deviations of replicate data. The proteome data obtained with the ISD and WinO methods were plotted using the UMAP (C). UMAP was performed using the umap package in R. Triangle and circle show the 100-cell and 10000-cell proteomics data, respectively. For details of sample preparation, see the legend of Figure 5.

Gene-ontology (GO) analysis of 798 proteins identified ribosomal proteins, proteasome-related proteins, and enzymes of the central carbon metabolism system (Figure 7A). In addition to these abundant proteins, transmembrane proteins (TMPs) and cell adhesion-related proteins were identified. To examine the effect of the WinO method on the recovery of TMPs, we compared the distribution of the number of transmembrane domains (TMDs) between 832 TMPs identified with the ISD method using 10000 cells with the 70 TMPs identified by the WinO method using 100 cells (Figure 7B). We found that the distribution patterns were highly correlated (*r* = 0.9826, *p* < 0.0001). TMPs are some of the most difficult proteins to identify using proteomics, and proteins with more TMDs are generally more difficult to extract and identify ^26,27^. The WinO method uses the PTS as protein extraction developed for membrane proteomics ^16,17^. These results indicated that the PTS improved the extraction and digestion efficiencies of membrane proteins, as well as soluble proteins. Thus, these results provided evidence that the WinO method-based proteomic analysis combined with PTS was an effective and unbiased approach for 100-cell proteomics.

**Figure 7.**
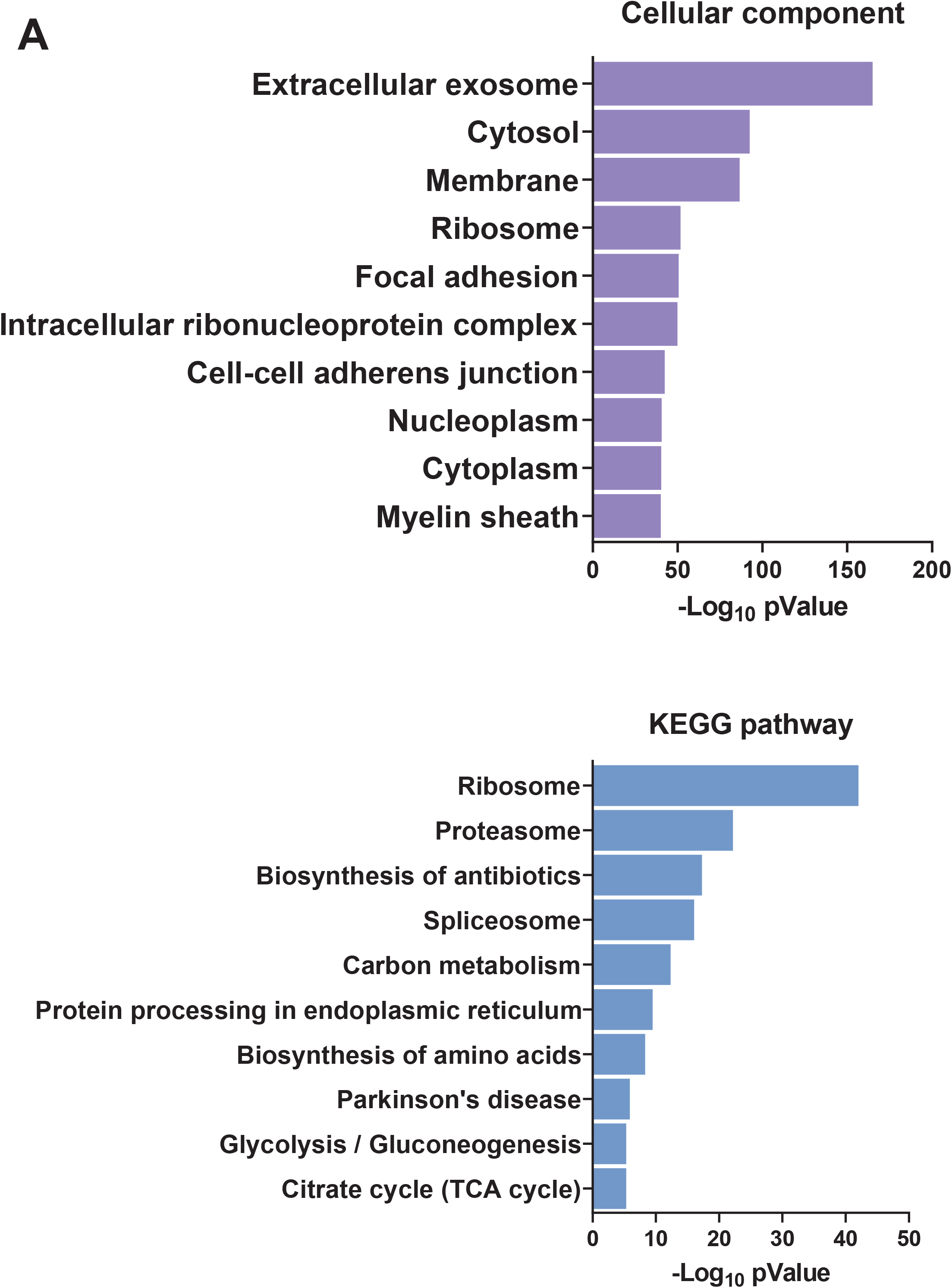
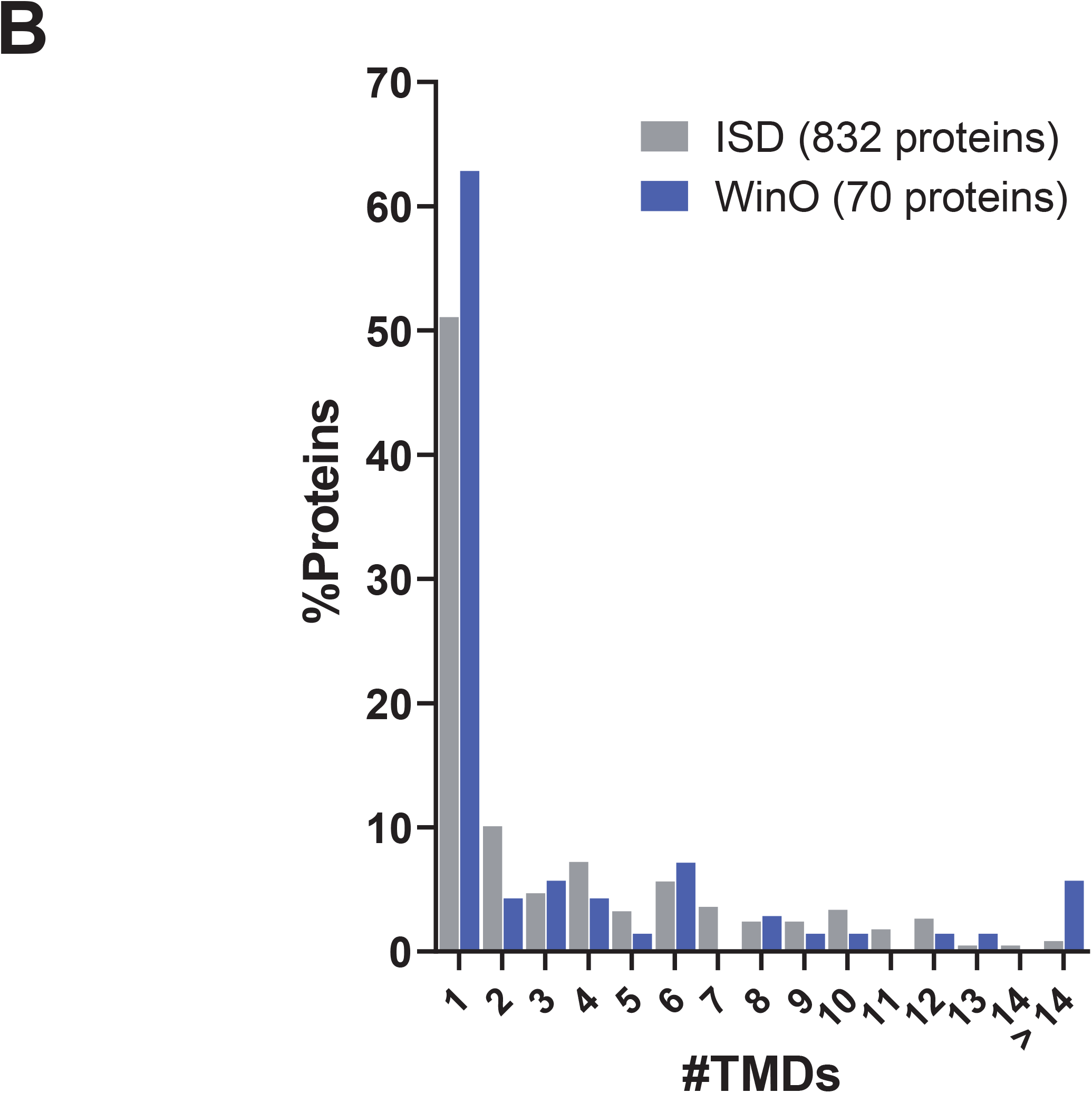
Protein identified by the WinO method. Gene ontology analysis with DAVID version 6.7 (https://david.ncifcrf.gov/summary.jsp) was performed using 798 proteins identified from 15 multiple myeloma cell lines by the WinO method (A). Distribution of transmembrane domains (TMDs) in proteins identified with either the ISD or WinO method (B). We identified 835 and 70 TM proteins from the 15 cell lines using the ISD and WinO methods, respectively. The number of TMDs in each protein was determined from the UniProt database (https://www.uniprot.org/).

### 4. Single-cell proteomic analysis using the WinO method

Finally, we examined the applicability of the WinO method for single-cell proteomics. Single RPMI8226 cells were directly sorted into a 96-well plate, and proteins were digested using the ISD or WinO method in quadruplicate. Peptides were labeled with TMT reagents and combined with TMT-labeled peptides that corresponded to 50 cells. The combined samples were then analyzed by nanoLC-MS/MS using an Orbitrap Eclipse, identifying 845 proteins and 2493 peptides. Of these identified proteins and peptides, 462 proteins (Table S9) and 1506 peptides (Table S10) were quantified. The average number of quantified peptides was 227.0 ± 114.5 and 1177.8 ± 131.6 for the ISD and WinO method, respectively (Figure 8A). The average number of quantified proteins was 140.8 ± 51.8 and 400.3 ± 32.5 for the ISD and WinO method, respectively (Figure 8A). The numbers of these peptides and proteins were significantly higher in the WinO method at 5.2-fold (*p* < 0.0001) and 2.8-fold (*p* < 0.0001), respectively (Figure 8A). Furthermore, to examine the effect of the WinO method on protein recovery, we compared the levels of proteins quantified by both methods. Figure 8B shows a volcano plot that compares 247 commonly identified proteins in both methods, indicating that the contents of 221 out of 247 proteins significantly (*p* < 0.05) increased 2-fold or more with the WinO method. There were no proteins significantly decreased in the WinO method. The median relative recovery of proteins was 10.21-fold greater with the WinO method than the ISD method. The levels of proteins commonly identified using both methods were significantly higher (*p* < 0.0001) than those uniquely identified using the WinO method (Figure 8C). These results suggested that the number of quantified proteins and peptides increased by increasing their recovery by the WinO method using single cells. In addition, 33 TMPs including one cluster of differentiation (CD) protein, CD71, were quantified in this study; among them, 24 TMPs were uniquely quantified using the WinO method. RapiGest ^13^ and n-dodecyl-β-D-maltoside ^14^ have been used for protein extraction in single-cell proteomics. It has been reported that these additives have comparable or higher solubility of membrane proteins than SDC ^17^, which was used in the WinO method. However, the Lys-C and trypsin activities were higher in the presence of SDC than these additives, resulting in a higher number of hydrophobic proteins and peptides identified ^17^. In this study, we used a mixture of SDC and SLS, which is known to considerably increase the solubility and the number of membrane as well as soluble proteins compared with SDC alone ^16^. These findings suggest that the extraction efficiency of proteins from a single cell is higher in the WinO method than in other single-cell proteomic techniques. Our WinO method enhanced protein recovery and protein identification not only soluble proteins but also TMPs from single cells, thereby highlighting its application for the single-cell proteomic analysis.

**Figure 8.**
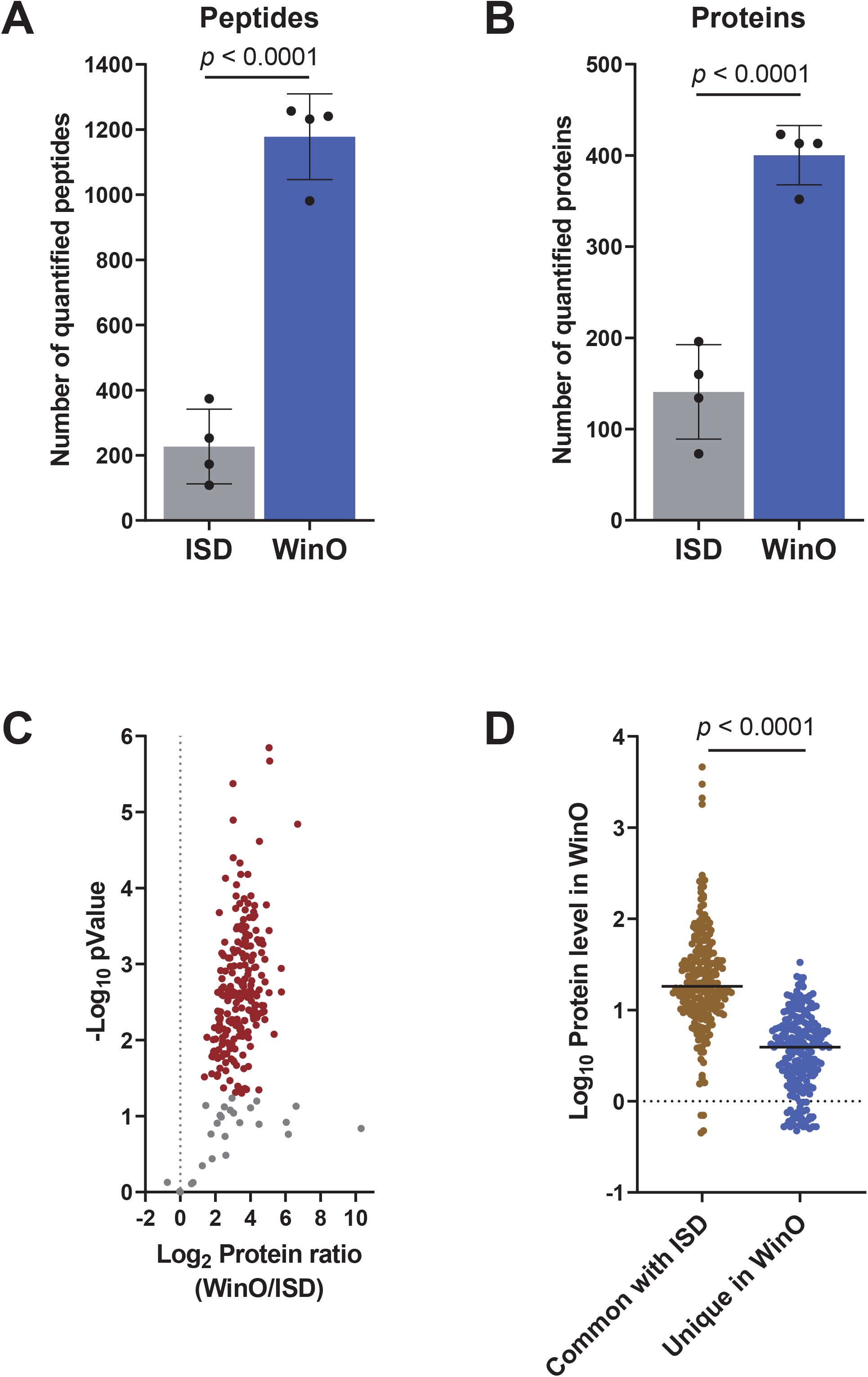
Application of the WinO method to single-cell proteomics. Numbers of proteins and peptides quantified using the ISD or WinO method (A). The graphs plot the average number of quantified proteins/peptides and the standard deviation of quadruplicate data. Protein levels detected from the WinO and ISD samples are compared in a volcano plot (B). Red dots indicate proteins with a significant (*p* < 0.05) change of 2-fold or more. (C) compares the Log10 protein levels obtained by the WinO method between commonly quantified with the ISD method (239 proteins) and the uniquely quantified in the WinO method (247 proteins). Each bar shows the median. Unpaired *t*-tests were performed using GraphPad Prism 8.4.3.

## Conclusions

On the WinO method, cells were directly injected into the sample droplet by a cell sorter. The method does not require any specialized equipment. The recovery of proteins and peptides is dramatically increased compared to the ISD method by reducing the contact area between the sample solution and the plastic container. In addition, the pipette tip does not contact the sample solution when the DTT, IAA, Lys-C, trypsin, and TMT solutions are added; thus, protein loss due to adsorption onto the pipette tip is avoided. Although there are still limitations to this method, such as the possibility of a lower peptide recovery rate once ethyl acetate is removed, the recovery of peptides and proteins increased approximately 10-fold for single-cell proteomics by coupling the use of phase transfer surfactants and carboxyl-coated hydrophobic beads. Several methods for single-cell proteomics have been previously reported 11,14,28. It is hardly possible to directly compare these methods in terms of numbers of quantified proteins due to differences in analytic systems and equipment. Nevertheless, we conclude that when compared to the ISD method, our novel strategy further improves the sensitivity of single-cell proteomics. In addition, although the WinO method was successfully performed on 96-well plates, we expect that the method is scalable to 384- and 1536-well plates using liquid handling robots, further enhancing the throughput of single-cell proteomics.

## Associated content

### Supporting Information

1. Supporting figures (PDF)

2. Table S1: Peptides identified from 100 cells by the ISD and WinO methods (XLSX)

3. Table S2: Proteins identified from 100 cells by the ISD and WinO methods (XLSX)

4. Table S3: Peptides identified from 100 cells using the WinO method with or without beads (XLSX)

5. Table S4: Proteins identified from 100 cells using the WinO method with or without beads (XLSX)

6. Table S5: Peptides identified from 100 cells by the WinO method (XLSX)

7. Table S6: Proteins identified from 10000 cells by the ISD method (XLSX)

8. Table S7: Proteins identified from 100 cells by the WinO method (XLSX)

9. Table S8: Proteins identified from 10000 cells by the ISD method (XLSX)

10. Table S9: Proteins identified from single cell by the ISD and WinO methods (XLSX)

11. Table S10: Peptides identified from single cell by the ISD and WinO methods (XLSX)

12. The MS raw data and result files have been deposited in the ProteomeXchange Consortium (http://www.proteomexchange.org/, PXD029814) via the jPOST partner repository (https://jpostdb.org, JPST001390) ^29^.

## Author Contributions

Conceptualization, T.M.; Investigation, T.M., Y.I., A.F., K.M., C.H.C., and D.K.; Formal analysis; T.M.; Resources, T.M., H.O., Y.K., and M.M.; Writing-original draft, T.M.; Funding acquisition, T.M., SE.O., and S.O.; Supervision, writing-review and editing, T.M., S.I., N.A., SE.O., and S.O.

## Notes

The Authors declare no competing financial interest.

## Acknowledgments

We thank Shio Watanabe, Kentaro Takahara, and Daisuke Higo from Thermo Fisher Scientific for conducting the single-cell proteomic analysis. This work was supported by JSPS KAKENHI Grant Numbers JP17K15042 and JP 19K05544 to T.M. This work was also supported by the Adaptable and Seamless Technology Transfer Program through Target-driven R&D (A-STEP) Grant Number JPMJTR20UM, and a Fusion Oriented Research for Disruptive Science and Technology Grant from Japan Science and Technology Agency (JST), and COCKPI-T Funding from Takeda Pharmaceutical Company Limited to T.M. This work was supported by JST-CREST (Grant Number JP171024167) to S.O. This work was also supported by grants from the National Institutes of Health issued under the award numbers R01AR065459 and R01GM129090 (S-E.O.)

