## Supplementary material for "Water droplet-in-oil digestion method for single-cell proteomics": Figure S

### Table of content

Figure S1 Comparison of the extraction efficiency with and without sonication and heating

Figure S2 Recovery of proteins and peptides in a water droplet in ethyl acetate

Figure S3 Comparison of fully cleaved peptide and mis-cleaved peptide recovery using ISD and WinO methods

Figure S4 Comparison of the addition of six types of magnetic beads on the recovery of peptides using the WinO method

Figure S5 Correlation of the recovery rate of peptides in the WinO method with beads and the frequency of basic or acidic amino acids

Figure S6 Distribution of %CV of the ISD and WinO methods

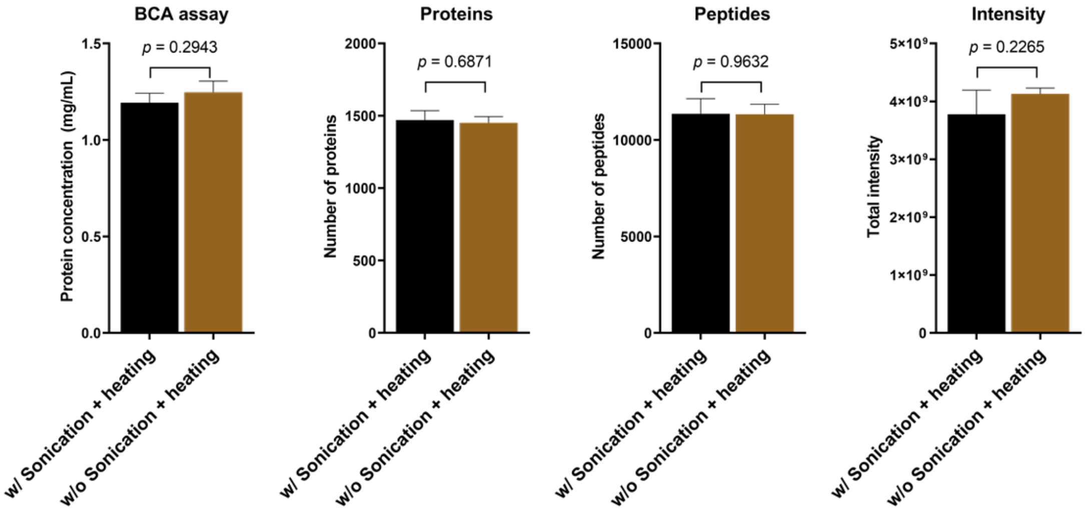


**Figure S1 Comparison of the extraction efficiency with and without sonication and heating**

The RPMI8226 cell pellet containing 5E5 cells were dissolved in 100 mL of 100 mM Tris-HCl (pH9.0) containing 125 unit of benzonase, 12 mM SDC, and 12 mM SLS. In one group, proteins were extracted by sonication for 20 min followed by heating at 95 °C for 5 min. In another group, proteins were extracted by leaving samples on the benchtop at 25 °C for 30 min. The extracted protein amounts were quantified by the BCA assay. We used 20 μL of the sample for protein digestion. These processes were performed in triplicate. The digested peptides were analyzed by nanoLC-MS/MS using TripleTOF 5600. Triplicate data were averaged, and error bars show standard deviation. Unpaired *t*-test was performed using GraphPad Prism 8.4.3.

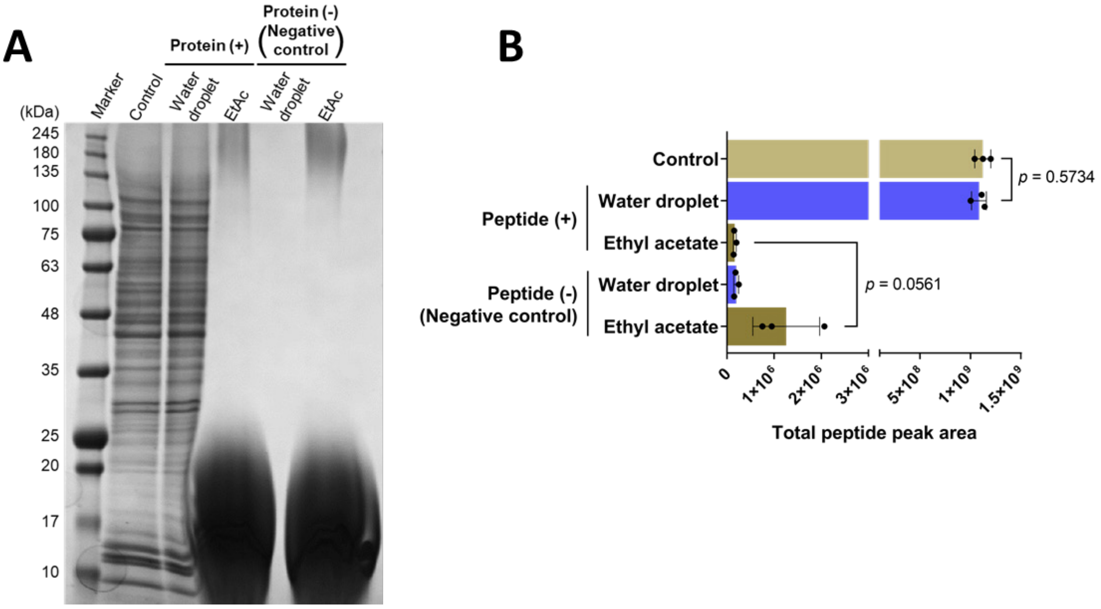


**Figure S2 Recovery of proteins and peptides in a water droplet in ethyl acetate**

To evaluate the protein and peptide recovery in a water droplet in ethyl acetate (EtAc), 10 µg HEK293 whole cell lysate or 10 µg digested peptides dissolved in 3 mM SDC, 3 mM SLS, 37.5 mM AmBic in 25 mM Tris-HCl (pH 9.0) were added into EtAc. After 24 h incubation, EtAc and water droplets were collected. The distributions of proteins or peptides in each fraction were confirmed by performing 5-20% SDS-PAGE (A) and nanoLC-MS/MS (B), respectively. As a control, protein or peptide solution used for this examination were also analyzed. Unpaired *t*-test was performed using GraphPad Prism 8.4.3.

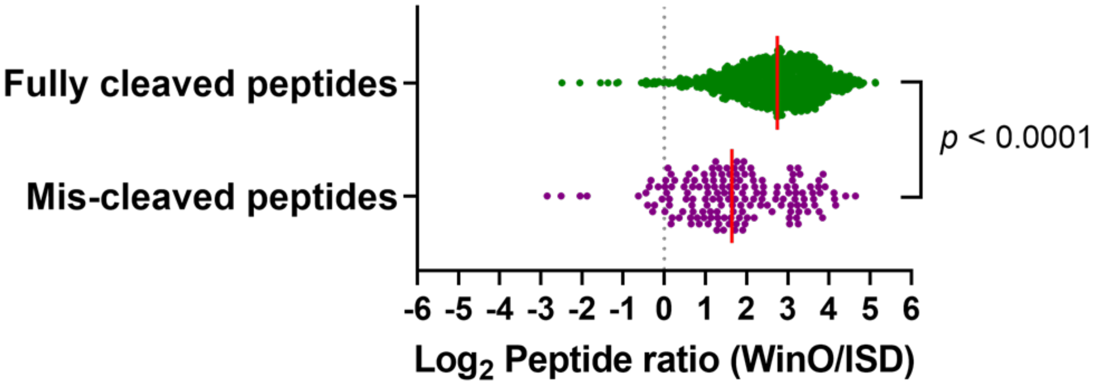


**Figure S3 Comparison of fully cleaved peptide and mis-cleaved peptide recovery using ISD and WinO methods**

The peptide recovery was calculated by dividing the peptide intensity obtained from the WinO method by that from the ISD method. Unpaired *t*-test was performed using GraphPad Prism 8.4.3. For details, see the legend of Figure 2.

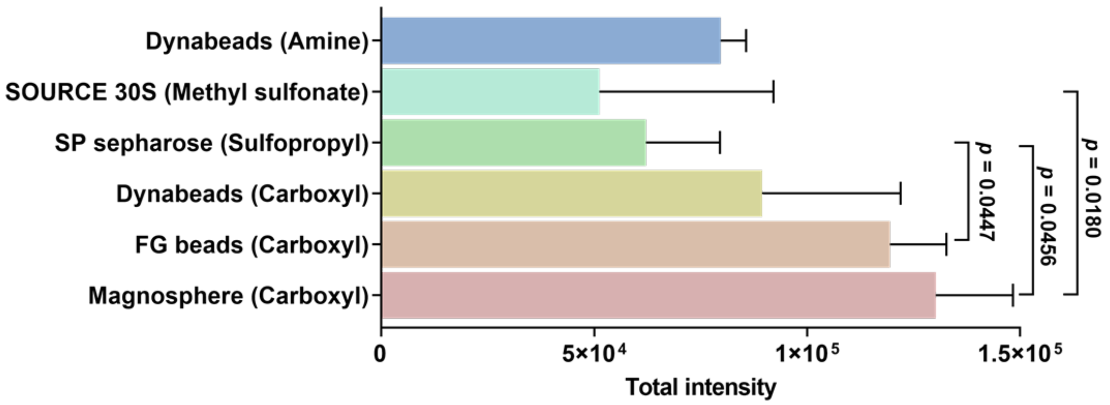


**Figure S4 Comparison of the addition of six types of magnetic beads on the recovery of peptides using the WinO method**

We evaluated the use of amine-coated Dynabeads, methyl sulfonate-coated SOURCE 30S beads, sulfopropyl-coated SP sepharose beads, carboxyl-coated Dynabeads, carboxyl-coated FG beads, or carboxyl-coated Magnosphere beads on the efficacy of the WinO method for protein and peptide preparation. The beads were equilibrated with 50 mM AmBic containing 12 mM SDC and 12 mM SLS. The WinO method was performed in triplicate using 10 ng HEK293 whole cell lysate in the presence of 1.65 mg beads. Each bar indicates the total intensity of the quantified peptides. Error bars show standard deviations. One-way ANOVA and multiple comparisons were performed using the GraphPad Prism 8.4.3. *p*-values showing the significant differences ($p < 0.05$) between samples are shown.

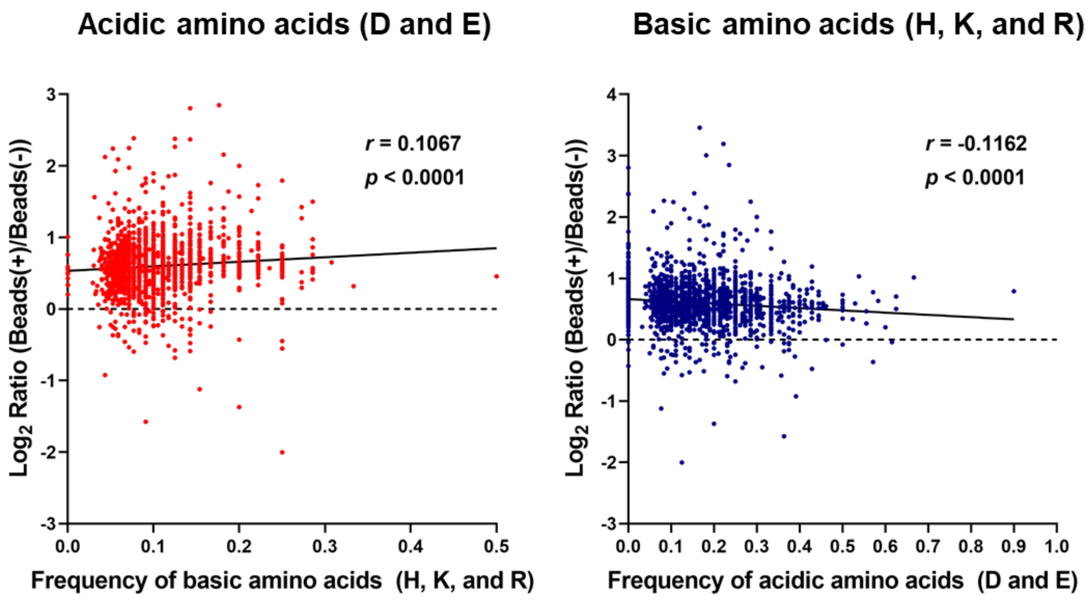


**Figure S5 Correlation of the recovery rate of peptides in the WinO method with beads and the frequency of basic or acidic amino acids**

The Pearson correlation and *p*-values were calculated using GraphPad Prism 8.4.3. For details, see the legend of Figure 4.

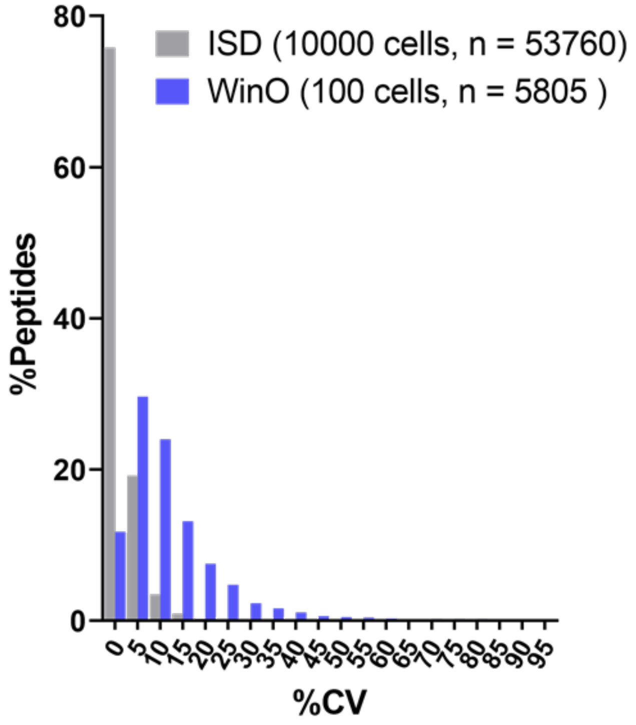


**Figure S6 Distribution of %CV of the ISD and WinO methods**

For the ISD and WinO methods, 10000 or 100 cells from 15 multiple myeloma cell lines were used for proteomics. The %CV of protein levels was calculated from data obtained from triplicate or quadruplicate analyses of each cell line. For details, see the legend of Figure 5.
